# Does Precision Grip Research Extend to Unconstrained, Multidigit Grasping?

**DOI:** 10.1101/2024.12.20.627508

**Authors:** Fabrizio Lepori, Frieder Hartmann, Kira Dehn, Manuela Chessa, Roland W Fleming, Guido Maiello

## Abstract

Most daily tasks require using our hands. Whether taking a sip from a glass or throwing a ball, we effortlessly select appropriate grasps. Yet, despite many possible hand configurations, most grasping research has focused on the finger-and-thumb ‘precision grip’. We thus questioned whether findings on precision grip hold under unconstrained grasping conditions. To test this, we compared how participants grasped 3D objects made of brass and wood, with both precision grip and unconstrained grasps. When unconstrained, participants rarely selected precision grips, favoring multi-digit grasps. Nevertheless, in both conditions, participants shifted grasps towards the objects’ center of mass and, when grasp factors conflicted, the variability in their selections increased, indicating greater uncertainty about the optimal strategy. Further, despite favoring multidigit grasps, participants consistently placed the thumb and index finger on the same positions on the objects, suggesting that in multidigit grasps, the additional fingers primarily provided support. Our findings thus reveal that object material affects unconstrained grasping similarly to precision grip and imply that previous precision grip research may extend to unconstrained, multidigit conditions.

**NEW & NOTEWORTHY:** Most grasping research focuses on two-digit ‘precision grips’, yet humans have more than two fingers. Here, we test whether previous precision grip findings apply to unconstrained grasping. We find that participants often use more than two digits when free to choose but consistently place thumb and index finger similarly on objects regardless of the number of fingers used. Our results thus highlight how the large body of precision grip literature can extend to multidigit grasping.

## INTRODUCTION

In everyday life we frequently grasp objects. This seemingly mundane action requires nuanced sensorimotor processes to select appropriate hand configurations to engage effectively with objects (1). Previous research has uncovered many of these processes, revealing that grasps are influenced by factors including an object’s material composition (2–5), shape (6–9), surface roughness (5, 10, 11), or orientation and position (5, 8, 12, 13). However, most of this prior research has focused on two-digit *precision grips*, and the few studies on multi-digit grasping focus on a single object (14) or very simple 2D shapes (3, 15). This is likely due to precision grip grasping being easier to investigate. For example, cables and markers affixed to fingertips can be used to record precision grip parameters such as the maximum grip aperture and the contact locations of the index and thumb fingertips on graspable objects (2, 5, 11, 16). Nevertheless, our everyday interactions with objects are not restricted to just index and thumb (17, 18). Indeed, attempts to classify human grasping have identified many distinct multidigit grasp types (17, 19). Does this mean that previous grasping literature is limited and misleading? To explore this, we investigated whether recent precision grip findings generalize to more natural, unconstrained and multidigit grasping behaviors.

Previously (2), we looked at how several factors interact in the selection of optimal, two-digit grasps on 3D objects made of various configurations of wood and brass cubes. We found that participants tended to shift their grasps towards the object’s center of mass. Here, we tested whether these previous findings on precision grip also apply to unconstrained grasping. We asked participants to grasp a subset of the objects from our previous study (2) in two separate sessions, while we tracked their hand movements. In one session, participants were given no specific instructions on how to interact with the object. In the other, they were instructed to perform two-digit precision grips, using exclusively thumb and index finger. We sought to test whether participants, when left unconstrained, would naturally gravitate towards using precision grip grasps. Our other objective was to test whether object mass and material configuration would affect unconstrained grasping similarly to precision grip.

## MATERIALS AND METHODS

### Participants

Twenty right-handed students/staff from Justus Liebig University Giessen (age range 20-39 years, mean 26.7, 15 female) participated for financial compensation (8 Euro/hour). Participants had normal or corrected-to-normal vision. Procedures were approved by Local Ethics Committee of Justus Liebig University Giessen and adhered to the principles outlined in the Sixth Revision of the Declaration of Helsinki (2008).

### Apparatus

Our setup is shown in **Figure 1**. Participants sat with their head in a chinrest 30 cm above a workbench. Target objects were positioned at a designated location on the workbench, 30 cm from the chinrest. We employed the motion tracking setup described in (20). The experiment, implemented in *Python 3*.*7*.*0*, employed state-of-the-art 3D motion tracking equipment and software (*Qualisys AB, Sweden*), which consisted of eight tracking (*Qualisys Miqus M5*) and six video cameras (*Qualisys Miqus Video*) arranged on a square frame surrounding the workspace (**Figure 1A**). To capture hand and object movements, we placed reflective markers at 24 specific locations on the back of participants’ hands (Figure 1B) and attached four uniquely configured markers to each stimulus object (Figure 1C).

**Figure 1.**
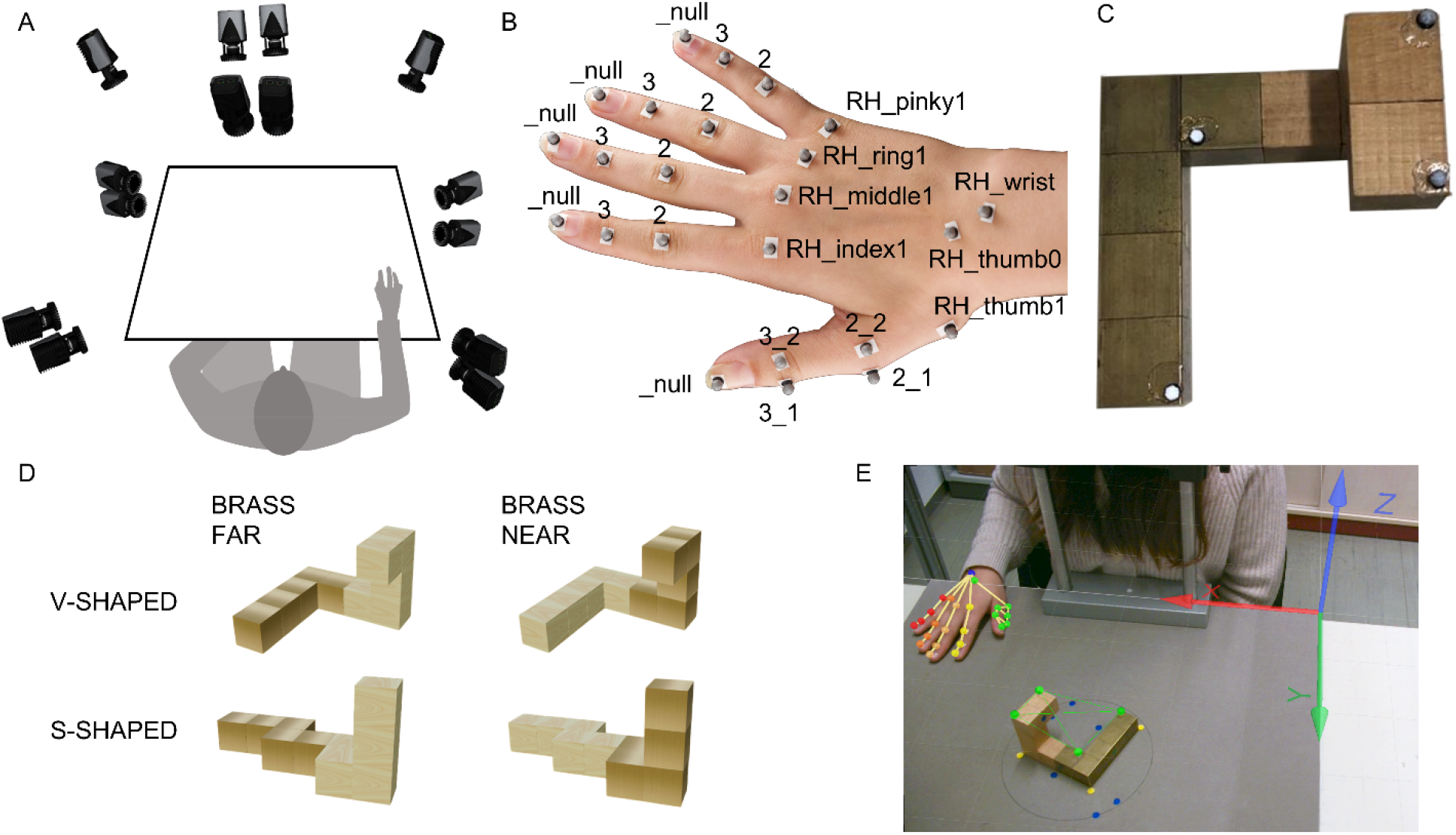
Experimental set-up. (**A**) Participants were seated at a workbench, imaged from all side by 14 cameras. (**B**) Markers were placed at specific landmarks on the back of a participant’s hand following (20). (**C**) One of four stimulus objects along with its unique markers configuration. (**D**) Virtual replicas of the four stimulus objects. (**E**) A still frame from one of the video cameras for the “V-shaped” object in the “brass far” configuration.

### Stimuli

A wooden parallelepiped object (7.5×2.5×2.5cm, 29.1g) was used as training stimulus. A subset of four different objects from (2), consisting of two distinct shapes in two material compositions, served as stimuli for experimental sessions. Each object was made of five wooden cubes (2.5 cm^3^, 9.7 g each) and five brass cubes (2.5 cm^3^, 133.5 g each). Total object weight was 716 g. The cubes’ arrangement formed a bipartite structure, with brass cubes connected to each other on one side and wooden cubes forming the opposing side. For same shaped objects, the materials arrangement was reversed, leading to distinct center of mass locations. For each stimulus object, we created a triangulated mesh replica (15380 vertices) using *Blender 3.1.2*. **Figure 1D** shows 3D renderings of all four stimulus objects in the orientation viewed by participants.

### Design and Procedure

The experiment consisted of a training session and two experimental sessions. In the training session, participants performed five trials, instructed to grasp the training object without constraints on the number of digits used. To avoid biasing participants towards precision grip grasping, the first experimental session was always unconstrained grasping, the second was precision grip grasping. In unconstrained grasping sessions, participants were asked to grasp the objects in a manner that felt most natural and comfortable to them, with no imposed limitation on the number of digits they could employ. In precision grip sessions participants were instead instructed to execute precision grips, using exclusively thumb and index finger to grasp stimulus objects.

Each session consisted of 20 trials, five repetitions for each of the four stimuli presented in randomized order. Prior to each trial participants were instructed to place their hand at a designated starting location on the workbench to the right of the chinrest (**Figure 1E**) and close their eyes. While participants’ eyes were closed, the experimenter positioned a stimulus at the target location on the workbench, using tape markers as reference. An auditory cue then signaled the start of a trial: participants then opened their eyes and reached out to grasp the object, lift it, place it back down, return their hand to the starting position, and close their eyes again. Participants had no time constraints, and the whole action took approximately 10 seconds.

## Data Analysis

Motion tracking data was recorded using *Qualisys Track Manager (QTM)* software and analyzed using *MATLAB R2023A*. Recorded data first underwent a pre-processing stage. For each participant a dedicated *AIM* (Automatic Identification of Markers) model was created using the relative positions of the markers on the participant’s hand, as depicted in **Figure 1B**. Each of the four objects employed in the experiment was instead identifiable by its unique marker configuration (**Figure 1E**). We calibrated the triangulated mesh replica for each object by defining the marker positions also on the mesh models and computing the appropriate rototranslation matrix.

## Time of first contact

In contrast to previous work (2, 5) using the method by Schot et al. (21), we adopted an alternative procedure to estimate the time of contact as it could not be easily adapted to unconstrained grasping. We identified the timepoint in which the velocity of the stimulus object along the vertical axis increased above a fixed threshold of 0.4 mm/s. We then selected the time of first contact as 150 ms (15 samples) prior to the object motion. We selected this delay using the precision grip sessions, by comparing the timepoint of object motion to the time of hand-object contact calculated using the Schot et al. (21) procedure. Across all participants and objects, the Schot et al. (21) procedure estimated the time of hand-object contact on average 15 samples earlier than the time of object motion (± 5 frames). We thus applied this average delay to both precision grip and unconstrained sessions.

## Key variables

Having identified the time of first contact, we next determined the *number of fingers in contact with the object*, and the corresponding *contact points* on the object. We did this by projecting the tracked fingertip positions onto the calibrated mesh model object replica. For each fingertip marker, we defined its *contact point* as the closest object vertex if their distance did not exceed a threshold of 25 mm (the approximate width of fingertip plus the marker size). if the distance was higher, we assumed that finger and object were not in contact. Having identified the contact points, we further defined the *grasp centroid* as the average location of the contact points involved in each grasp. From this, we computed each grasp’s *signed distance from the object centroid* as the Euclidean distance between the grasp centroid and the object centroid, where grasps farther than the centroid were defined as positive, and distances closer than the centroid were defined as negative. For each participant in each condition, we also computed the *grasp variability* as the standard deviation of the signed distance from the object centroid across the five trial repetitions. Finally, we estimated *the contact time* for each digit as the time when the digit reached minimum distance from the object, relative to the time of object motion.

All estimated contact points on the objects were manually inspected by the experimenter. In rare instances (7% of trials) the procedure produced implausible contact points (e.g., thumb and index contact points both on the same object surface). These instances were reprocessed by the experimenter, who manually selected the time of first contact by watching the video recordings alongside the motion tracking data. Very few trials (2%) were excluded for other reasons (e.g., participant mistakes, tracking occlusions).

### Statistical Analyses

We used a two-by-two (brass far vs brass near; unconstrained vs precision) within-subject analysis of variance (ANOVA) to analyze *number of fingers used, grasp centroid distance, and grasp variability data*. Wilcoxon signed-rank tests were used to compare contact times for different digits. *P-values* < .05 were considered statistically significant.

## RESULTS

Twenty participants grasped wood and brass objects in two configurations, with the brass side near or far from the participant’s right-hand starting position. Participants grasped these objects in two different sessions, with no constraints on the hand configuration (unconstrained session, **Figure 2A**) or using only thumb and index finger (precision grip session, **Figure 2B**).

**Figure 2.**
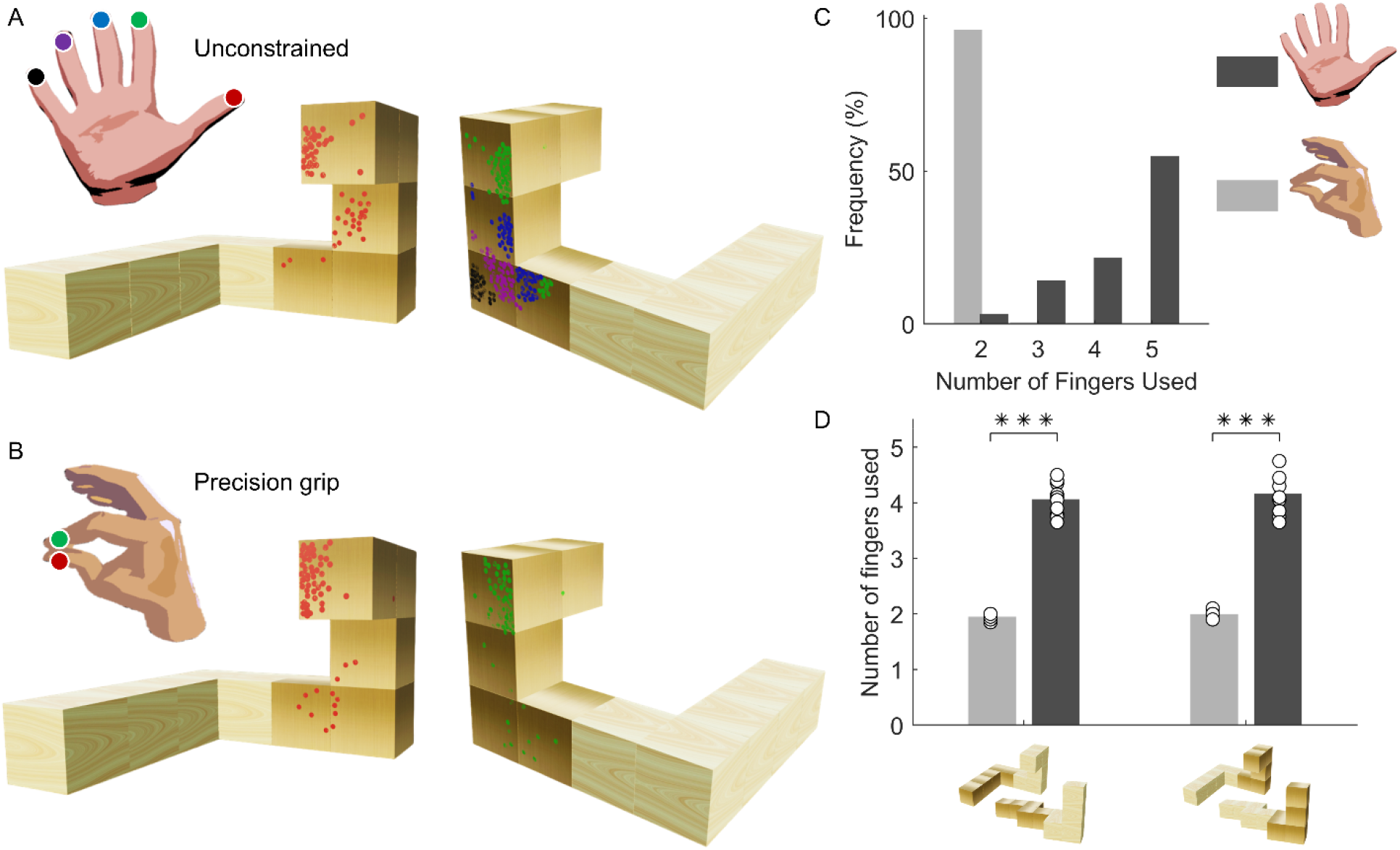
Participants used different number of digits in precision grip and unconstrained grasping sessions. (**A**,**B**) Contact points onto one stimulus object during unconstrained grasping (**A**) and precision grip (**B**) sessions. Objects are shown from the same (left) and opposite side (right) presented to participants. Dots are individual participant grasps. Different colors correspond to different digits. (**C**) Frequency of number of digits employed in precision and unconstrained grasping sessions across all participants and objects. (**D**) Average number of fingers used in precision and unconstrained grasping sessions for brass far (left) and brass near (right) objects. Bars are means; circles are data for individual participants. *** p < .001

### Participants did not spontaneously employ precision grip

To test whether participants spontaneously employed precision grip, we compared the number of digits used in unconstrained and precision grip sessions (**Figure 2C**). As instructed, participants used only two digits in precision grip sessions^1^. In unconstrained sessions instead, participants almost never selected only two digits, and in over 50% of cases used all five digits. ANOVA on the number of digits used (**Figure 2D**) revealed a significant main effect of grasp type (*F*(1,19) = 196.90, *p* < .001), no significant effect of material composition (*F*(1,19) = 1.16, *p* = .296), and no significant interaction, (*F*(1,19) = 0.18, *p* = .676).

## Material composition affects the selection of grasping points

We next assessed the influence of material composition on the selection of hand-object contact points. **Figure 3** shows two interesting patterns. In the brass near condition for both precision and multidigit sessions, (**Figure 3A,B**, right) grasps all clustered on the brass side of the object. In the brass far conditions (**Figure 3A,B**, left), grasps were more scattered and fell on both wood and brass sides of the object.

**Figure 3.**
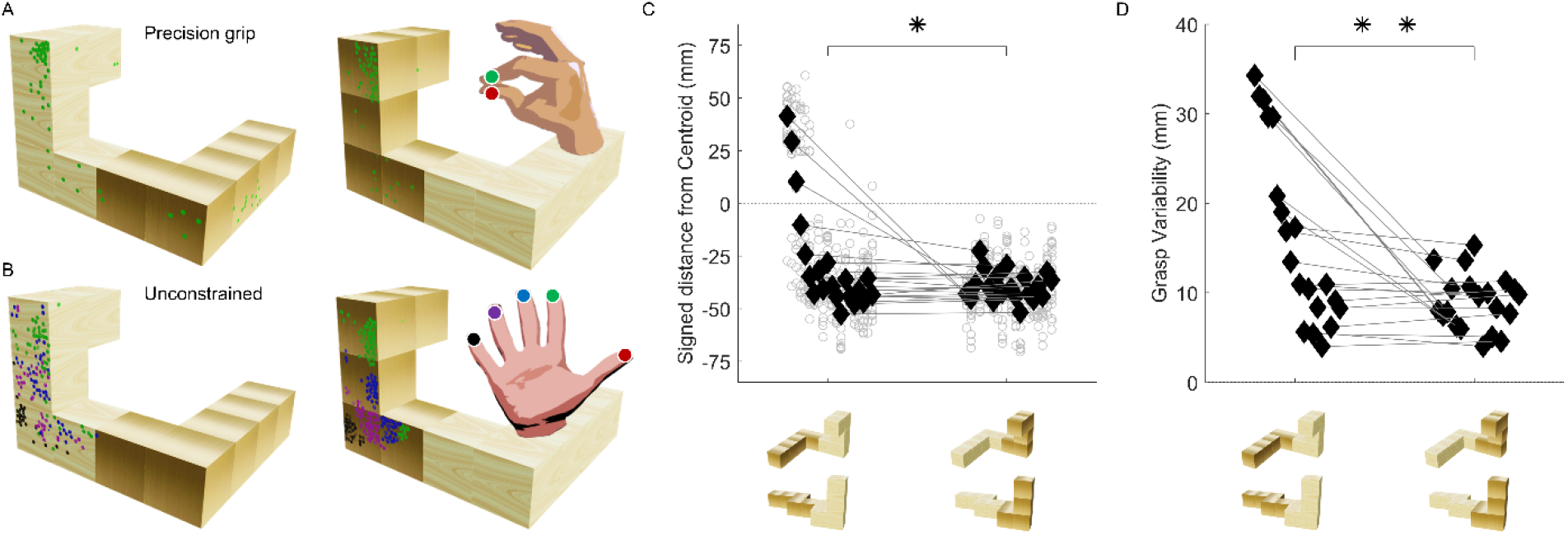
Effects of object material configuration. (**A**,**B**) Contact points onto two stimulus objects of the same shape but different material configuration during precision grip (**A**) and unconstrained (**B**) grasping sessions. Objects are viewed from the opposite side presented to participants to show contact locations of the digits opposing the thumb. Dots are individual grasps. Different colors correspond to different digits. (**C**) Signed distance from the object centroid for brass near (right) and brass far (left) material configurations. Light grey circles are individual trials, thin lines connect individual participant medians (black diamonds) in brass far and brass near configurations. The dotted line at 0 represents the position of the object’s centroid. Negative values indicate that the contact occurred on the side of the object closer to the participant’s right hand starting position; positive values indicate contact on the farther side. (**D**) Grasp variability for brass near (right) and brass far (left) material configurations. Thin lines connect individual participant’s grasp variability (black diamonds) in brass far and brass near configurations. In C and D, datapoints are ordered by the size of the material effect. * *p* < .05, ** *p* < .01

**Figure 3C** shows that in the brass near condition, participants uniformly grasped the object on the heavier brass side, which was also the side nearest to their right-hand starting position. In the brass far condition instead, a few participants switched completely and grasped the object on the side farthest from their hand starting position, and most of the rest who kept grasping on the wood side still shifted their grasps by a few millimeters towards the brass side. ANOVA on the signed distance between grasp and object centroids confirmed a significant main effect of material composition (*F*(1, 19) = 5.19, *p* = .034), no significant effect of grasp type (*F*(1,19) = 0.25, *p* = .621) and no significant interaction (*F*(1, 19) = 1.69, *p* = .209). Thus overall, the grasp centroid location was affected by the object’s material composition but was not significantly affected by the grasp technique.

**Figure 3D** shows that grasp variability was significantly lower in the brass near condition compared to the brass far condition (main effect of material composition, *F*(1,19) = 9.31, *p* = .007), and there was no significant effect of grasp type (*F*(1,19) = 1.56, *p* = .228) and no significant interaction (*F*(1,19) = 2.18, *p* = .156). This suggests that in the brass far condition, in which different grasp factors were in conflict (2, 4, 11), in both unconstrained and precision grip conditions participants were uncertain about where they should grasp the object and across repeated trials they experimented with different grasp locations.

### The index finger sets the grasp in motion

Finally, we assessed the contact time of each digit, in relation to the time of first object motion, in precision and unconstrained grasping sessions independently of objects and material composition (**Figure 4A,B**). We found that the thumb was consistently the last finger to touch the object in both precision (*z* = 2.05, *p* = .040) and multidigit sessions (*z* = 3.92, *p* < .001). In unconstrained sessions, the middle, ring, and pinky fingers reached the objects first. However, both in precision and unconstrained grasping, the objects began to move only once the index finger was in contact, suggesting that the index plays a critical role in initiating object movement across both grasp types.

**Figure 4.**
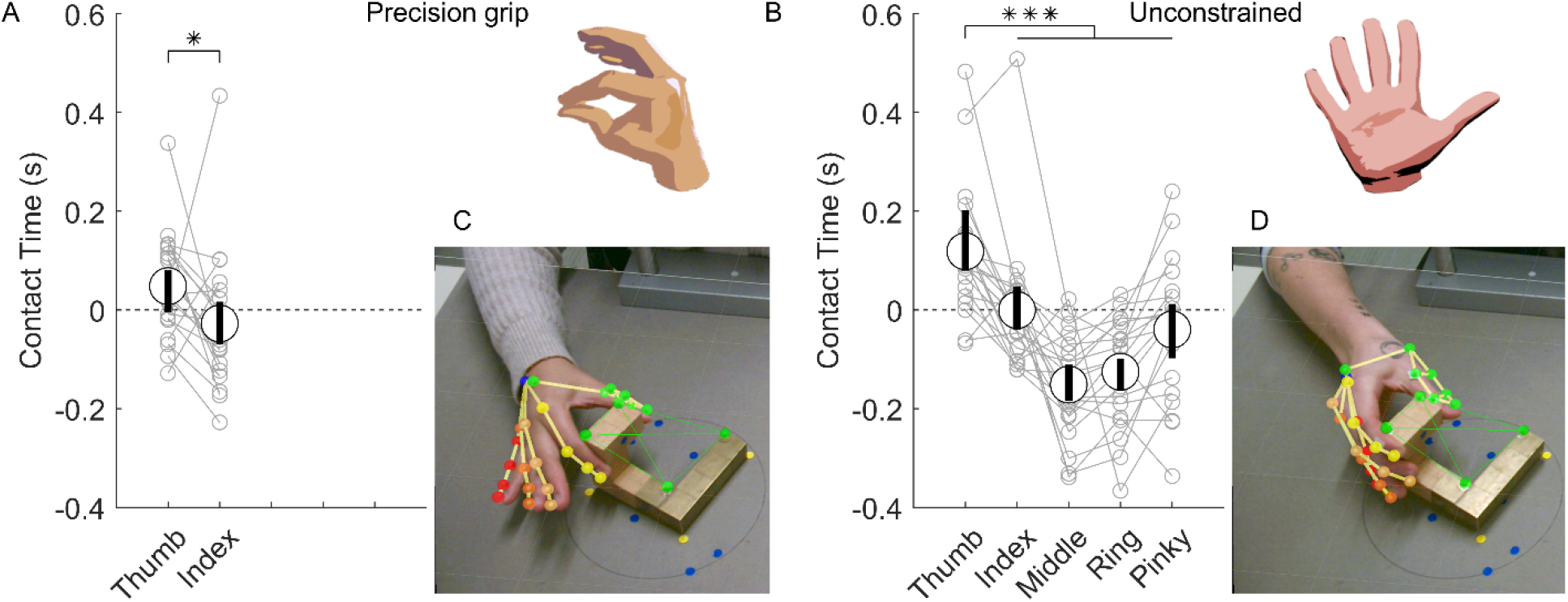
Are additional digits primarily for support? (**A**,**B**) Contact times for each digit in precision grip (**A**) and unconstrained (**B**) grasping sessions. Thin connected markers are data from individual participants. Large markers are means across participants, error bars are within-subject 95% confidence intervals. (**C**,**D**) Side-by-Side single frame of two participants grasping the brass-far object in precision grip (**C**) and unconstrained sessions (**D**).

## DISCUSSION

We find that, for the objects tested here, precision grip was not participants’ preferred grasping technique when they were free to choose. Instead, they favored multi-digit grasps. Nevertheless, participants had no difficulties in employing only two digits when explicitly instructed to do so. This agrees with findings from Feix et al. (22) who proposed that although multidigit grasps are ideal for heavier objects, they are extremely versatile and can be effective also in cases when a precision grip would suffice. Does this mean that precision grip studies (6–8, 23, 24) do not generalize to unconstrained grasping situations?

Our results suggest they do. Our findings replicate and extend previous research on the impact of material composition on grasping behavior. For example, Paulun et al. (5) showed that participants grasp single-material objects near their center of mass if the material is heavier, while Klein et al. (2) found that participants adjust their grasps for multi-material objects by shifting the grasp location according to the object’s center of mass. Here, we replicated a subset of the conditions from Klein et al. (2) with both precision grip and multi-digit grasping. Our findings from the precision grip session reproduce the influence of the object’s material composition found in existing literature (2, 5). Importantly, we found this effect also generalized to the unconstrained sessions. This implies that material composition influences human behavior regardless of the grasping technique employed.

Previous work also shows humans prefer grasps that optimize object visibility by grasping objects from the side nearest to their hand (2, 24, 25). Here, participants also consistently grasped objects from the side closest to them when the heavier side was near. When the heavier side was farther instead, grasping objects from nearer side conflicted with the tendency to grasp close to the center of mass to avoid high grip torques (2, 23, 26). This conflict likely introduced uncertainty, prompting participants to experiment with various grasping locations. Such behavior suggests that in scenarios where object properties do not yield a single, optimal grasping area, humans adapt by sampling the various options.

We note two limitations of the current study. First, although the effect of material composition was reliable, it was more modest than anticipated based on the overall results from Klein et al. (2). Additionally, precision grip sessions always occurred after unconstrained grasping sessions. This was purposeful to avoid biasing participants to select precision grip grasps in the unconstrained sessions. Nevertheless, it is in principle possible that session order could have biased precision grip selections towards unconstrained locations. Both these potential issues are resolved however by noting that our precision grip results closely match the data from the exact subset of conditions replicated from the Klein et al. (2) study (c.f. supplementary figure S1 from Klein et al. (2)). This suggests that object material configuration had similar effects across studies, and that participants’ precision grip grasps were largely unaffected by prior unconstrained grasps. Nevertheless, extending the current work to the selection of hand-object contact surfaces (27) could reveal additional, nuanced differences between precision grip and unconstrained grasping (28).

To summarize, when free to use any number of digits, participants rarely chose only two, yet the effect of object material was the same independently of the number of digits employed. This raises the question, if the use of thumb and index finger remains consistent across grasp configurations (even in terms of relative timing), what are the remaining three fingers for? Visual inspection of the executed grasps gives us a clue (**Figure 4C,D**). Participants appeared to select similar thumb and index contact points in both unconstrained and precision grip sessions. The main difference across sessions was that the additional fingers in unconstrained grasping sessions provided additional support. We speculate that the additional fingers employed in unconstrained conditions served primarily to reinforce and stabilize the grasp. This observation is good news for previous literature, as it suggests that precision grip findings extend to unconstrained grasping.

## DATA AVAILABILITY

Data and analysis scripts will be made available from the Zenodo data repository (doi upon final publication).

## GRANTS

This research was supported by:

The DFG (SFB-TRR-135-TP-C1: “Cardinal Mechanisms of Perception”, and project PA 3723/1-1).

Research Cluster “The Adaptive Mind” funded by the Excellence Program of the Hessian Ministry of Higher Education, Science, Research and Art.

An ERC Advanced Grant (ERC-2022-AdG-101098225 “STUFF”) to RWF.

A Mayflower PhD Scholarship awarded to FL.

## AUTHOR CONTRIBUTIONS

FL, FH, KD, MC, RF, GM conceived and designed the research. FL, FH, KD performed the experiments. FL and GM analyzed the data. FL, FH, KD, MC, RF, GM interpreted the results of the experiments. FL and GM prepared the figures. FL and GM drafted the manuscript. FL, FH, KD, MC, RF, GM edited and revised manuscript. FL, FH, KD, MC, RF, GM approved final version of manuscript.

## Footnotes

1 The frequency of two-digit precision grip grasps is < 100 % because in a small percentage of precision grip trials (3.25 %) our data processing algorithm identified only one contact point on the object.

## REFERENCES

1. Napier JR. The prehensile movements of the human hand. The Journal of Bone and Joint Surgery British volume 38-B: 902–913, 1956. doi: 10.1302/0301-620X.38B4.902.

2. Klein LK, Maiello G, Paulun VC, Fleming RW. Predicting precision grip grasp locations on three-dimensional objects. PLoS Comput Biol 16: e1008081, 2020. doi: 10.1371/journal.pcbi.1008081.

3. Lukos J, Ansuini C, Santello M. Choice of Contact Points during Multidigit Grasping: Effect of Predictability of Object Center of Mass Location. Journal of Neuroscience 27: 3894–3903, 2007. doi: 10.1523/JNEUROSCI.4693-06.2007.

4. Maiello G, Schepko M, Klein LK, Paulun VC, Fleming RW. Humans Can Visually Judge Grasp Quality and Refine Their Judgments Through Visual and Haptic Feedback. Front Neurosci 14: 591898, 2021. doi: 10.3389/fnins.2020.591898.

5. Paulun VC, Gegenfurtner KR, Goodale MA, Fleming RW. Effects of material properties and object orientation on precision grip kinematics. Experimental Brain Research 234: 2253–2265, 2016. doi: 10.1007/s00221-016-4631-7.

6. Chen Z, Saunders JA. Online processing of shape information for control of grasping. Experimental Brain Research 233: 3109–3124, 2015. doi: 10.1007/s00221-015-4380-z.

7. Eloka O, Franz VH. Effects of object shape on the visual guidance of action. Vision Research 51: 925–931, 2011. doi: 10.1016/j.visres.2011.02.002.

8. Lederman SJ, Wing AM. Perceptual judgement, grasp point selection and object symmetry. Experimental Brain Research 152: 156–165, 2003. doi: 10.1007/s00221-003-1522-5.

9. Schettino LF, Adamovich SV, Poizner H. Effects of object shape and visual feedback on hand configuration during grasping. Experimental Brain Research 151: 158–166, 2003. doi: 10.1007/s00221-003-1435-3.

10. Burstedt MK, Flanagan JR, Johansson RS. Control of grasp stability in humans under different frictional conditions during multidigit manipulation. J Neurophysiol 82: 2393–2405, 1999. doi: 10.1152/jn.1999.82.5.2393.

11. Klein LK, Maiello G, Fleming RW, Voudouris D. Friction is preferred over grasp configuration in precision grip grasping. Journal of Neurophysiology 125: 1330–1338, 2021. doi: 10.1152/jn.00021.2021.

12. Christopoulos VN, Schrater PR. Grasping Objects with Environmentally Induced Position Uncertainty. PLoS Computational Biology 5: e1000538, 2009. doi: 10.1371/journal.pcbi.1000538.

13. Mamassian P. Prehension of objects oriented in three-dimensional space: Experimental Brain Research 114: 235–245, 1997. doi: 10.1007/PL00005632.

14. Crajé C, Lukos JR, Ansuini C, Gordon AM, Santello M. The effects of task and content on digit placement on a bottle. Exp Brain Res 212: 119–124, 2011. doi: 10.1007/s00221-011-2704-1.

15. Gilster R, Hesse C, Deubel H. Contact points during multidigit grasping of geometric objects. Experimental Brain Research 217: 137–151, 2012. doi: 10.1007/s00221-011-2980-9.

16. Derzsi Z, Volcic R. MOTOM toolbox: MOtion Tracking via Optotrak and Matlab. Journal of Neuroscience Methods 308: 129–134, 2018. doi: 10.1016/j.jneumeth.2018.07.007.

17. Feix T, Romero J, Schmiedmayer H-B, Dollar AM, Kragic D. The GRASP Taxonomy of Human Grasp Types. IEEE Transactions on Human-Machine Systems 46: 66–77, 2016. doi: 10.1109/THMS.2015.2470657.

18. Stival F, Michieletto S, Cognolato M, Pagello E, Müller H, Atzori M. A quantitative taxonomy of human hand grasps. J NeuroEngineering Rehabil 16: 28, 2019. doi: 10.1186/s12984-019-0488-x.

19. Naceri A, Moscatelli A, Haschke R, Ritter H, Santello M, Ernst MO. Multidigit force control during unconstrained grasping in response to object perturbations. Journal of Neurophysiology 117: 2025–2036, 2017. doi: 10.1152/jn.00546.2016.

20. Hartmann F, Maiello G, Rothkopf CA, Fleming RW. Estimation of Contact Regions Between Hands and Objects During Human Multi-Digit Grasping. JoVE : 64877, 2023. doi: 10.3791/64877.

21. Schot WD, Brenner E, Smeets JBJ. Robust movement segmentation by combining multiple sources of information. Journal of Neuroscience Methods 187: 147–155, 2010. doi: 10.1016/j.jneumeth.2010.01.004.

22. Feix T, Bullock IM, Dollar AM. Analysis of Human Grasping Behavior: Correlating Tasks, Objects and Grasps. IEEE Trans Haptics 7: 430–441, 2014. doi: 10.1109/TOH.2014.2326867.

23. Kleinholdermann U, Franz VH, Gegenfurtner KR. Human grasp point selection. Journal of Vision 13: 23–23, 2013. doi: 10.1167/13.8.23.

24. Paulun VC, Kleinholdermann U, Gegenfurtner KR, Smeets JBJ, Brenner E. Center or side: biases in selecting grasp points on small bars. Experimental Brain Research 232: 2061–2072, 2014. doi: 10.1007/s00221-014-3895-z.

25. Maiello G, Paulun VC, Klein LK, Fleming RW. Object Visibility, Not Energy Expenditure, Accounts For Spatial Biases in Human Grasp Selection. i-Perception 10: 204166951982760, 2019. doi: 10.1177/2041669519827608.

26. Biegstraaten M, Smeets JBJ, Brenner E. The relation between force and movement when grasping an object with a precision grip. Exp Brain Res 171: 347–357, 2006. doi: 10.1007/s00221-005-0271-z.

27. Moscatelli A, Bianchi M, Serio A, Terekhov A, Hayward V, Ernst MO, Bicchi A. The Change in Fingertip Contact Area as a Novel Proprioceptive Cue. Current Biology 26: 1159–1163, 2016. doi: 10.1016/j.cub.2016.02.052.

28. Adams MJ, Johnson SA, Lefèvre P, Lévesque V, Hayward V, André T, Thonnard J-L. Finger pad friction and its role in grip and touch. J R Soc Interface 10: 20120467, 2013. doi: 10.1098/rsif.2012.0467.

